# Detection and Classification of Cardiac Arrhythmias by a Challenge-Best Deep Learning Neural Network Model

**DOI:** 10.1101/766022

**Authors:** Tsai-Min Chen, Chih-Han Huang, Edward S. C. Shih, Yu-Feng Hu, Ming-Jing Hwang

**Affiliations:** Institute of Biomedical Sciences, Academia Sinica, Taipei, Taiwan; Genome and Systems Biology Degree Program, Academia Sinica and National Taiwan University, Taipei, Taiwan; Division of Cardiology, Department of Medicine, Taipei Veterans General Hospital, Taipei, Taiwan; Institute of Clinical Medicine and Cardiovascular Research Institute, National Yang-Ming University, Taipei, Taiwan.

## Abstract

**Background:** Electrocardiogram (ECG) is widely used to detect cardiac arrhythmia (CA) and heart diseases. The development of deep learning modeling tools and publicly available large ECG data in recent years has made accurate machine diagnosis of CA an attractive task to showcase the power of artificial intelligence (AI) in clinical applications.

**Methods and Findings:** We have developed a convolution neural network (CNN)-based model to detect and classify nine types of heart rhythms using a large 12-lead ECG dataset (6877 recordings) provided by the China Physiological Signal Challenge (CPSC) 2018. Our model achieved a median overall F1-score of 0.84 for the 9-type classification on CPSC2018’s hidden test set (2954 ECG recordings), which ranked first in this latest AI competition of ECG-based CA diagnosis challenge. Further analysis showed that concurrent CAs observed in the same patient were adequately predicted for the 476 patients diagnosed with multiple CA types in the dataset. Analysis also showed that the performances of using only single lead data were only slightly worse than using the full 12 lead data, with leads aVR and V1 being the most prominent. These results are extensively discussed in the context of their agreement with and relevance to clinical observations.

**Conclusions:** An AI model for automatic CA diagnosis achieving state-of-the-art accuracy was developed as the result of a community-based AI challenge advocating open-source research. In-depth analysis further reveals the model’s ability for concurrent CA diagnosis and potential use of certain single leads such as aVR in clinical applications.

**Abbreviations:** CA, cardiac arrhythmia; AF, Atrial fibrillation; I-AVB, first-degree atrioventricular block; LBBB, left bundle branch block; RBBB, right bundle branch block; PAC, premature atrial contraction; PVC, premature ventricular contraction; STD, ST-segment depression; STE, ST-segment elevation.

## Introduction

Cardiac arrhythmias (CAs) are harbingers of cardiovascular diseases and could cause deaths [1]. CAs are diagnosed by electrocardiogram (ECG), a non-invasive, inexpensive and widely applied clinical method to monitor heart activities. To diagnose CAs, the wave-like features known as P wave, QRS wave, T wave, etc. of ECG are examined. A complete ECG usually contains recordings from six limb leads (I, II, III, aVR, aVL, aVF) and six chest leads (V1, V2, V3, V4, V5, V6), with each lead measuring electrical activity from a different angle of the heart on both the vertical plane (for limb leads) and the horizontal plane (for chest leads) [2, 3].

These different leads exhibit distinct features of ECG signals associated with specific types of CA. The following are some examples. Atrial fibrillation (AF) is characterized by the fibrillatory atrial waves and irregular conduction of QRS [4, 5]. Left bundle branch block (LBBB) is diagnosed by the distinct QRS morphology at leads I, aVL, V1, V2, V5, and V6, while right bundle branch block (RBBB) is diagnosed by the rsR’ pattern at V1 and V2 [6]. First-degree atrioventricular block (I-AVB) is defined as constant PR intervals longer than 0.2 second [7]. The premature atrial contraction (PAC) and premature ventricular contraction (PVC) indicate the electrical impulse from an abnormal site: Namely, the P wave or QRS morphology of PAC and PVC is different from those in normal heart beats [8, 9]. ST segment is abnormal if ST-segment elevation (STE) is greater than 0.1 mV or ST-segment depression (STD) is greater than 0.1 mV [10].

To reliably recognize these complex CA-associated ECG characteristics, considerable training is required. Indeed, studies have shown that internists or cardiologists sometimes misdiagnosed CA types [11, 12]. The significant growth of ECG examination which increases physician’s workload and burnout aggravates the problem. This problem can be alleviated by developing computer algorithms to assist the physician with accurate and automatic diagnosis. Although such a task is difficult owing to the large variance in the geometrical and physiological features of ECG signals [13], significant progress has been made, especially in recent years [14].

There are generally two approaches to develop an automatic CA diagnostic tool. The first one would split ECG signals into the units of the heartbeat, or cycles of the characteristic ECG waveforms. Thus, even with a small number of subjects, this beat-based approach can generate a large amount of beat data for machine learning to train predictive classification models. However, extracting ECG morphological features to delineate ECG signals proves challenging, as it is often an imprecise undertaking [14]. And while prediction accuracies as high as >99% have been reported in beat-based studies, they could be masked by the fact that both training and test beats can come from the same individual. As a result in one study, when test beats were taken from patients not included in the training set, the cross validation accuracy of a six types CA classification decreased from 99.7% to 81.5% [15].

The MIT-BIH Arrhythmia Database (MIT-BIH AD) [16, 17] and the UCI Machine Learning Repository: Arrhythmia Data Set (UCIAD) [18], which respectively contain only 48 and 452 subjects, have been the source of publicly available ECG data for most of previous CA prediction studies. However, databases of a small number of subjects such as these two would tend to cause over-fitting problems for classification, especially for neural network algorithms [19]. Data over-fitting would also arise from significantly unbalanced data, i.e., data being unproportionally concentrated in one or few CA types. These are problems that can produce biased results when analyzing MIT-BIH AD and UCIAD [20, 21]. For instance, in a study analyzing UCIAD, a high accuracy (92%) of CA classification was achieved when data were split into 80% in the training set and 20% in the test set, but the accuracy dropped to only 60% when the training-test splitting was 50-50 [20]. Additional drawbacks of using the two databases are that ECG data only included two leads (e.g. leads II and V1, II and V5, II and V4, and V2 and V4) in MIT-BIH AD, and only extracted features (average width of Q, amplitude of Q, etc.) but not the raw data of 12-lead ECG are available in UCIAD.

The second approach provides an end-to-end solution, avoiding the main difficulty of the beat-based approach. This requires a very large ECG database as well as the construction of a suitable deep learning artificial neural network to take advantage of the large database. Developments on both aspects in recent years have made the second approach increasingly attractive. For example, to promote open-source research, the PhysioNet/Computing in Cardiology Challenge 2017 (CinC2017) released single-lead (lead I) ECG data of 8,528 subjects with four labeled CA types (AF, normal, other rhythms, noise) to the public [22]. Using convolutional neural network (CNN) plus 3 layers of long short-term memory (LSTM, one kind of recurrent neural network (RNN)), Xiong et al. produced the top performance of CinC2017 with an F1 score (the harmonic mean of the precision and recall) of 0.82 on its hidden test set (3,658 subjects) [23].

As CinC2017, The China Physiological Signal Challenge 2018 (CPSC2018) hosted by the 7th International Conference on Biomedical Engineering and Biotechnology [24] released a large ECG database for free download and set aside a hidden test set to assess models submitted by challenge participants from around the world. Different from CinC2017, the ECG data of CPSC2018 were 12–lead and subjects were grouped into normal and eight types of CA: AF, I- AVB, LBBB, RBBB, PAC, PVC, STD, and STE. This represents the biggest 12–lead ECG database with the most labeled CA types in the public domain to date. Here, we report a deep learning artificial neural network modeling of the CPSC2018 ECG data, and the results that won the first place in the competition.

## Methods

The CPSC2018 ECG database has been described in detail by Liu and coworkers [24]. Briefly, a total of 9831 12–lead ECG recordings from 9458 individuals were collected from eleven hospitals in China. The ECG was sampled by a frequency of 500 Hertz for a few seconds to a minute, with a few exceptions including one lasting as long as 144 seconds. Each recording was also labeled as the normal type or eight abnormal CA types as mentioned above. The database was divided by a random 70-30 training-test split, and only the training set was made available to the public. Gender and age distribution between the training set and the test set were fairly balanced, so were the distributions of the subjects from the eleven hospitals and the CA types [25]. Of the 6877 training-set recordings, 470 received two CA-type labels and 6 received three.

Our model was built on a combined architecture of five CNN blocks, followed by a bidirectional gated recurrent unit (GRU), an attention layer [26, 27], and finally a dense, i.e. fully connected, layer (Fig 1). Within each CNN block there were two convolution layers and they were followed by a pooling layer to reduce the amount of parameters and computation in the network and control over-fitting [28]. Furthermore, between these CNN blocks or between other independent layers, including the one between the last CNN block and the bidirectional GRU layer, we randomly dropped 20% of their connections. We chose to use CNN and RNN because of their demonstrated ability to handle noisy signals and time series data in studies which included ECG classification [29, 30]. GRU is a new form of RNN proposed recently that can require less training time and less number of iterations than LSTM [31, 32]. We used batch normalization to adjust and scale the input from the attention layer, which determines a vector of importance weights, to the dense layer [33]. LeakyReLU activation function, a leaky version of Rectified Linear Unit, was used for each layer, except for the dense layer, where Sigmoid activation function was used [34].

**Fig 1.**
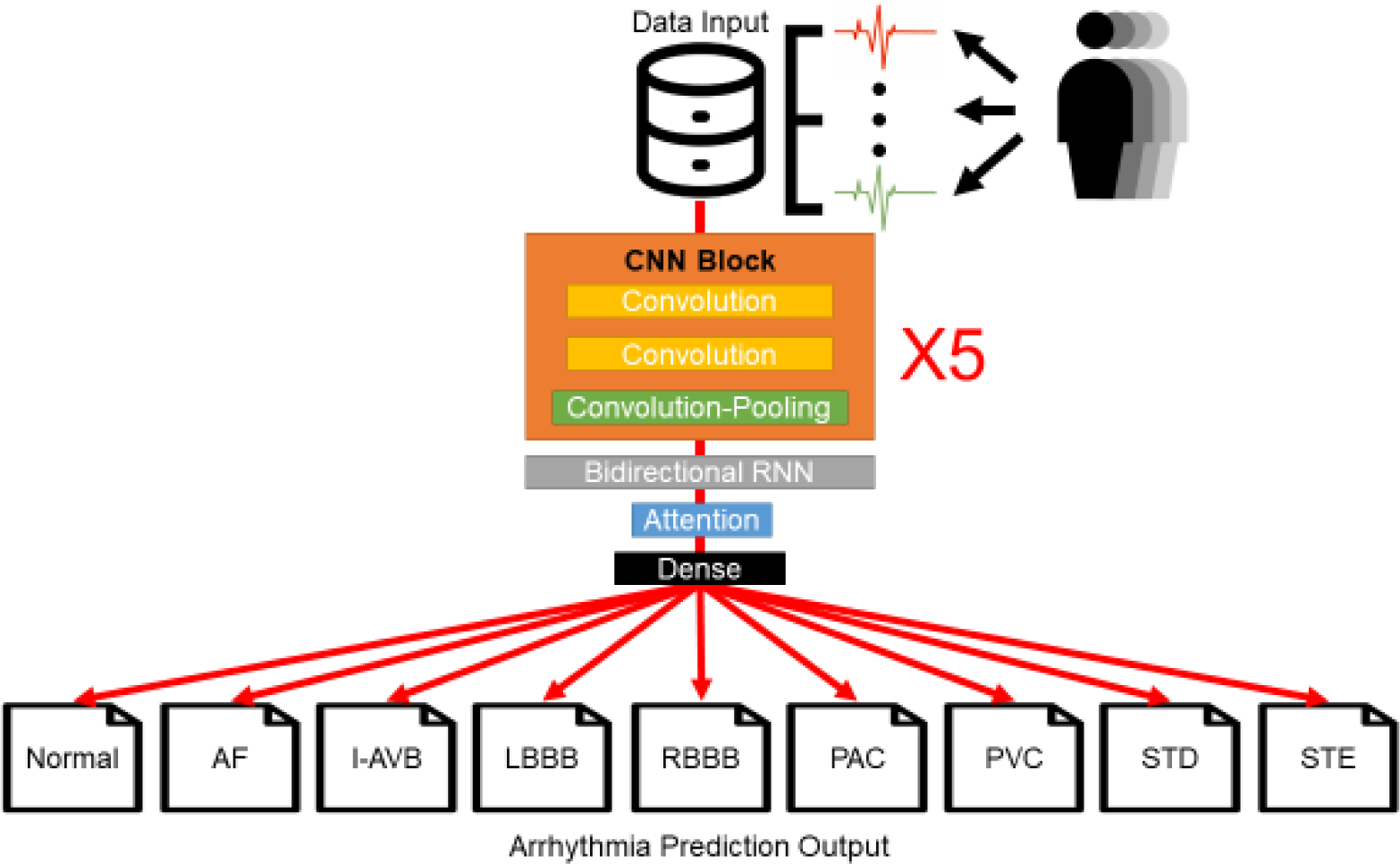
The architecture of deep learning artificial neural network for 12-lead ECG CA detection and classification. Layers and blocks are specified in rectangle boxes; “X5” indicates that five convolution neural network (CNN) blocks are tandem-connected before connecting to the bidirectional recurrent neural network (RNN) layer, which is a GRU layer. The output layer at bottom contains the probabilities predicted by the model for each of the nine types of the CA classification. The type with the highest probability is the type predicted by the model for the input ECG recording.

In our implementation, the CPSC2018 ECG data were processed into a matrix of three elements: the first is the subject ID, the second identifies which of the ECG’s 12 leads being considered, and the third contains its 72,000 ECG values, which correspond to the recordings taken by the maximum recording time (144 seconds) and on a frequency of 500 Hertz. We padded zeroes up front for any recording that was less than the maximum time. The 476 multi-labeled subjects were extracted when the rest of 6,401 subjects were randomly divided into 10 equal parts to set up an 8-1-1 train, validation and test scheme of machine learning. The extracted multi-labeled subjects were then added back to be included for the training. Our classification training was carried out using categorical-cross-entropy loss function and ADAM optimizer in the GPU version of TensorFlow from the Keras package [35-37]. Models were evaluated on their performance on the validation set for 100 training epochs (an epoch refers to one cycle through the full training dataset in artificial neural network learning). The best model, the one with the smallest loss on the validation set, was further evaluated by computing its F1-score on the test set. The procedure was repeated 10 times to complete the 10-fold training and validation plus test to produce 10 best validation models. The median F1-score for each CA label, including the normal type, for the 10 test sets was calculated using the F1-score package from Scikit-learn [38].

We further investigated the performance of using only single lead data. To do that, for a given lead we simply assigned zero to all the ECG values of the other 11 leads and derived the model using the same network architecture and the same 10-fold cross validation plus test procedure described above. This resulted in 120 best single-lead validation models and a median F1-score for each of the 12 single leads on each of the nine CA labels.

To compete for CPSC2018, the 130 best validation models (10 from full-lead training and 120 from single-lead training) were combined into one ensemble model by which the average of the output probabilities from the 130 models for each CA type was adjusted by a weight vector to produce the final probability for that CA type. The weights of the vector were optimized by genetic algorithm [39] to produce the best overall median F1-score on the 10 test sets. Given an input of an ECG recording, the CA type receiving the largest probability from the ensemble model would then be the type of CA predicted for that ECG recording. The ensemble model was our model submitted to CPSC2018, its performances on the hidden test set (2954 recordings) as computed and reported by CPSC2018 organizers are presented in Table 1.

**Table 1:** Comparison of model performances on tests^*^.

| CA Type | Best Validation Models |  |  | Ensemble Model |  |  |  |
| --- | --- | --- | --- | --- | --- | --- | --- |
|  | Median Accuracy | Median AUC (95% CI) | Median F1-score | Median Accuracy | Median AUC (95% CI) | Median F1-score | **Hidden Set F1-score** |
| Normal | 0.940 | 0.890<br>(0.810-0.942) | 0.795 | 0.949 | 0.867<br>(0.832-0.973) | 0.808 | **0.801** |
| AF | 0.969 | 0.928<br>(0.902-0.985) | 0.897 | 0.983 | 0.963<br>(0.914-0.993) | 0.944 | **0.933** |
| I-AVB | 0.972 | 0.899<br>(0.864-0.988) | 0.865 | 0.977 | 0.950<br>(0.875-0.990) | 0.899 | **0.875** |
| LBBB | 0.990 | 0.914<br>(0.748-1.000) | 0.821 | 0.995 | 0.942<br>(0.763-1.000) | 0.899 | **0.884** |
| RBBB | 0.955 | 0.956<br>(0.887-0.988) | 0.911 | 0.952 | 0.946<br>(0.871-0.976) | 0.903 | **0.910** |
| PAC | 0.957 | 0.867<br>(0.749-0.955) | 0.734 | 0.963 | 0.920<br>(0.779-0.981) | 0.797 | **0.826** |
| PVC | 0.970 | 0.928<br>(0.841-0.988) | 0.852 | 0.977 | 0.932<br>(0.864-0.996) | 0.874 | **0.869** |
| STD | 0.951 | 0.878<br>(0.797-0.972) | 0.788 | 0.959 | 0.906<br>(0.815-0.970) | 0.834 | **0.811** |
| STE | 0.976 | 0.707<br>(0.558-0.995) | 0.509 | 0.977 | 0.773<br>(0.603-0.993) | 0.600 | **0.624** |

\*These are results of the best validation models and the ensemble model on the ten 10-fold tests, except for those in the last column (boldfaced), which are the ensemble model's median F1-scores for the hidden test set of CPSC2018 reported at its website http://2018.icbeb.org/Challenge.html, which did not provide accuracy and AUC results.

## Results

### (1) Best validation models on 10-fold tests and ensemble model on hidden test

In Table 1, for each CA type the median accuracy, AUC (area under the receiver operating characteristic curve) and F1-score for the ten 10-fold tests from the best validation models are compared with those of the ensemble model, as well as with the F1-score of the ensemble model on the hidden test set of CPSC2018. The comparisons show that the ensemble model performed somewhat better than the best validation models, which is expected because the former combined and optimized the latter to produce the best 10-fold test results (see Methods). In addition, the ensemble model’s performance was quite stable across all CA types going from the publicly available data to the hidden test data, reflecting the fairly similar compositions of the two sets of data, as mentioned above.

Table 1 also reveals differential difficulties in predicting these CA types. Namely, the prediction accuracy decreased from AF, bundle branch blocks, premature contractions to ST abnormalities, with the normal type being one of the more difficult-to-predict types. The model’s prediction for STE had the lowest F1-score (0.5∼0.6), which may due in part to physician’s variable opinions on how to diagnose STE [40]. The same trend, including the prediction of the normal type, was observed in all other top-performing models of CPSC2018 (S1 Table). Indeed, almost all the top models produced very high F1-scores (> 0.9) for AF and bundle branch blocks. Our model had significantly better predictions than the other models on several CA types, especially PAC, PVC, STD, and STE. This explained how we outperformed others (S1 Table). It should be noted that all top models performed well (overall F1-score > 0.8) and the difference between our model and the second-place model was minimal (S1 Table).

### (2) Concurrent CA types

One reason for models to perform less accurately on certain CA types is that for some patients multiple CA types are predicted with almost equal probabilities. Fig 2 displays the probabilities output by the best validation models for ECG subjects when they were in the test fold of the10-fold tests. As may be seen, Normal, STD and STE are three types lacking a probability score that can make them stand out from the other eight types, in consistence with the model’s performance results presented in Table 1. Further analysis on model probabilities showed that for many AF patients, a common concurrent CA was RBBB, while many RBBB patients were often concurrent with PAC and PVC, in addition to AF (Fig 2). These probability results of concurrent CAs agreed well with the statistics of the 476 multi-labeled subjects: Namely, the three most multi-labeled incidences in these subjects are AF/RBBB, RBBB/PAC and RBBB/PVC (Table 2). An ensemble model without these 476 multi-labeled subjects being added back to the training set (see Materials and Methods) performed well in predicting these multiple CA labels (S3 Table and S4 Table), indicating the model’s ability to capture ECG features of concurrent CAs. These results are also generally compatible with clinical observations that rate-dependent (phase 3) block during ectopic atrial beats or AF could lead to RBBB [41] [42]. However, a larger dataset of multi-labeled subjects is required to fully evaluate our model’s performance on concurrent CA diagnosis.

**Fig 2.**
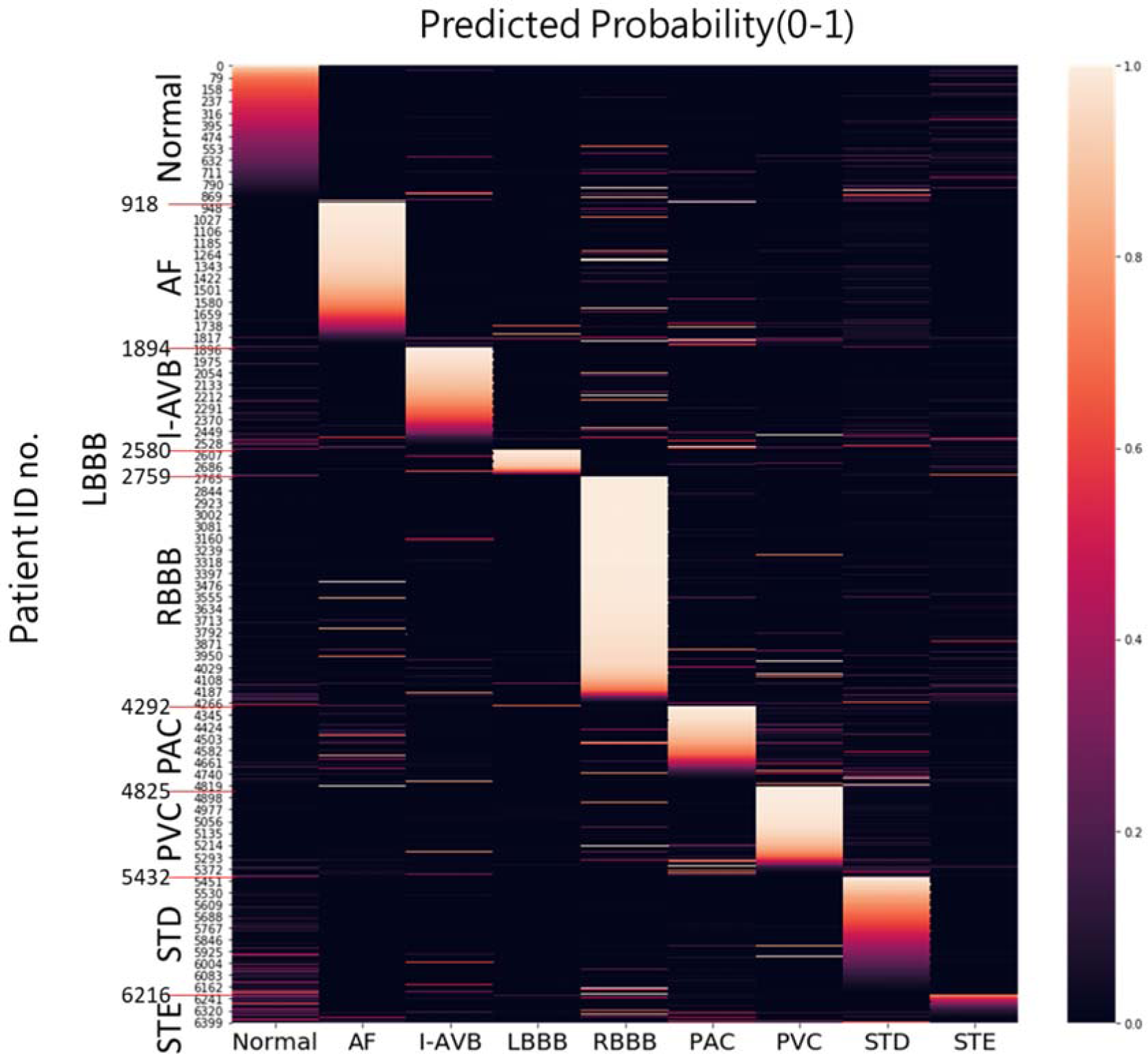
Probabilities output by the best validation models in the test fold of the 10-fold test. On the right is the color-coded probability scale.

**Table 2:** Label count statistics of the 476 multi-labeled subjects in the released CPSC2018 dataset^*^.

|  | AF | I-AVB | LBBB | RBBB | PAC | PVC | STD | STE |
| --- | --- | --- | --- | --- | --- | --- | --- | --- |
| AF | 0 | 0 | 29 | **172** | 4 | 8 | 33 | 2 |
| I-AVB |  | 0 | 8 | 10 | 3 | 5 | 6 | 4 |
| LBBB |  |  | 0 | 0 | 10 | 6 | 3 | 4 |
| RBBB |  |  |  | 0 | **55** | **51** | 20 | 19 |
| PAC |  |  |  |  | 2 | 3 | 6 | 5 |
| PVC |  |  |  |  |  | 0 | 18 | 2 |
| STD |  |  |  |  |  |  | 0 | 2 |
| STE |  |  |  |  |  |  |  | 0 |

\* Only the upper triangle portion of the symmetrical concurrent CA label counts is shown. The three largest counts are boldfaced.

### (3) Model performances with single lead

The median F1-scores for models of a single lead on the 10-fold tests are presented in Fig 3. The performances for the best validation models using the 12-lead data in Table 1 were largely replicated by those using only single-lead data. In most cases, only minimal changes of F1-scores for the classification of individual CA types were noted between the analysis of 12-lead and single-lead ECGs. The results also indicate aVR was one of the best-performing single leads, as its performance ranked first in overall average and three individual CA types (Normal, AF and STD), and also within top 3 in all CA types except STE and PAC. Another well-performing single lead is lead V1, which ranked first in three types (I-AVB, RBBB and PAC), but did worse than most other leads in some types. In comparison, lead I, which was used by Apple Watch [43], wasn’t as remarkable in our tests. Lead II, the favorite of the 12 leads by physicians to take a quick look at an ECG recording due to its clearest signal [44], ranked fifth in the overall average but was statistically no different from the leading leads (p value of paired t-test < 0.05). These results are largely supported by Bayes factor analysis [45] to rigorously assess statistical differences between these leads (see S5 Table, S6 Table, S7 Table).

**Fig 3.**
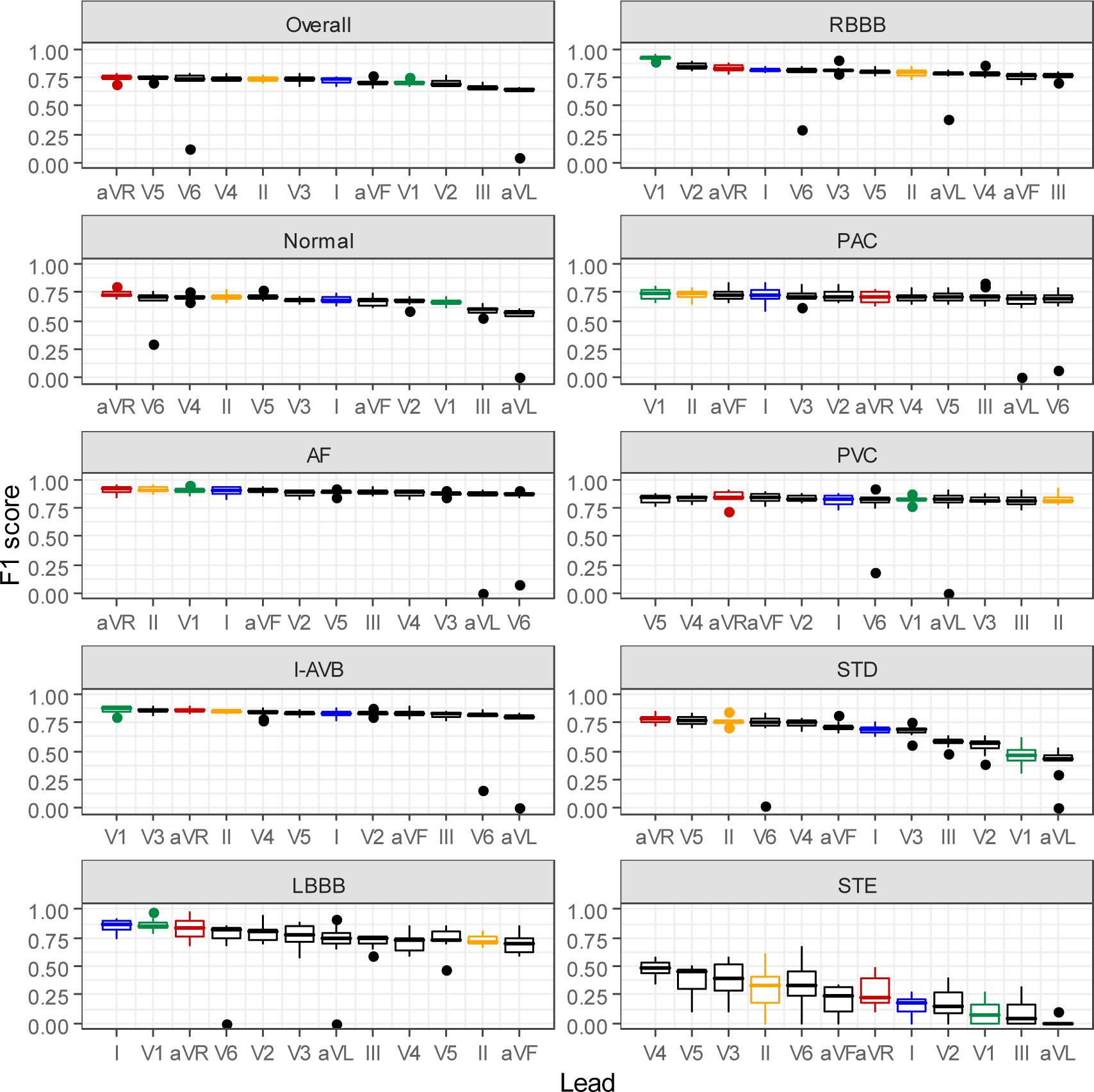
The ranked F1-score results of single lead models. The F1-scores (on the y axis) are from the single lead models performed on the 10-fold tests (see Materials and Methods). Lead aVR is shown in red, V1 in green, I in blue, and II in orange.

These performance rankings suggest the current model identified the lead-specific morphology of the CA types. For examples, the deep and broad S-waves in lead V1 and the broad clumsy R-waves in V6 had been used for the diagnosis of LBBB [46], and V1 and V6 were identified among the single leads with the leading performance. Meanwhile, the diagnosis criteria of RBBB included the rSR’ pattern in leads V1 and V2 [47], which were also selected as top–performing single leads.

## Discussion

Recent years have witnessed a number of successful applications using deep learning of artificial intelligence (AI) to make medical diagnosis [48]. The present work for CA detection and classification is related to the competition in CinC2017 [22] and studies that had been published recently [11, 22]. A direct performance comparison for the different studies is difficult because not all of them used publicly available ECG data and different CA types and type numbers had been predicted. The complexities of these deep learning models were also different: e.g., the total number of neural network layers is 18 in our model, comparing to 33 [11] and 5∼7 [22] in others. Nevertheless, all these studies seemed to achieve an overall F1-score around 0.82∼0.84. Although not fully tested in the real-world scenario, AI-based ECG diagnosis has been shown to significantly improve diagnosis accuracy, compared to general physicians and cardiologists [11, 12] (also see S2 Table for a very small sampling). Therefore, these AI models are capable of reducing erroneous diagnoses and medical overload. While this is very encouraging, it is a sobering reminder that until most of the “ground truth” diagnoses used to derive AI models are made by expert cardiologists, there might be a limit to how much model accuracy can be further improved.

Our analyses suggest that models built on single-lead information could predict CA types with minimal difference of performance from the 12 leads. The clinical diagnostic criteria of CA types are often lead-specific. The top-ranking single lead for RBBB or LBBB in our model was compatible with the leads in the diagnostic criteria of RBBB and LBBB [6], solidifying the validity of the present AI diagnosis model. The performance of aVR, a frequently clinically-ignored lead, in our AI model is intriguing and deserves attention. The leads I, II, and V1 are conventionally used as the modified leads in continuous monitoring or mobile device of ECG [43, 49]. In our AI model, aVR could predict a variety of CA types with a better performance than these conventional leads. The vector of lead aVR is parallel to the anatomical and corresponding electrical axis from atrial base to ventricular apex, and thus may maximize the electrical signals of atrial and ventricular depolarization. In comparison, lead I, which is used in Apple Watch for AF detection [50], did not perform as well in our analysis. Our results suggest the best predictive single lead for different CA types could be different for clinical applications. Our results may provide an impetus for future studies to investigate the potential use of lead aVR in different CA types and ECG devices (wearable or portable).

CAs are complex and concurrent CA types are not uncommon, especially for those that are related in cardiac electrophysiology. Although ECG-based CA diagnosis models have so far focused only on single-type predictions, our analysis shows that AI is capable of multi-type CA diagnosis. Detection and classification of concurrent CAs should be a subject for future studies and our model is a first step in that direction.

ECG has been shown capable of disease/health detection beyond CA, including, for example, the prediction of asymptomatic left ventricular dysfunction [51] and non-invasive potassium tracking [52]. As methods of AI machine learning continue to be advanced and made friendlier for non-AI specialists to employ, we can expect ECG to be explored for its diagnostic power in many more diseases and clinical applications.

## Conclusion

We developed a deep learning AI model capable of cardiologist-level CA detection and classification. The model was derived from a very large 12-lead ECG dataset made available for free access in a challenge competition to promote open-source research. Besides achieving the first placebest performance in the competition, the model was shown to yield promising results for two aspects worthy of future investigations in the field: concurrent CA diagnosis and use of less attended single leads such as aVR in clinical applications.

## Acknowledgements

We thankfully acknowledge the CPSC Challenge Chair Prof. Chengyu Liu and his conference coworkers for their help and making the ECG data publicly available.

## Data Availability Statement

All relevant data are at http://2018.icbeb.org/Challenge.html.

## Funding

The research of Hwang lab is supported by Institute of Biomedical Sciences, Academia Sinica, and Ministry of Science and Technology, Taiwan, grant number MOST108-2311-B-001-017.

## Competing interests

All authors have declared that no competing interests exist

## Author Contributions

Conceptualization: Tsai-Min Chen, Chih-Han Huang, Edward S. C. Shih, Ming-Jing Hwang.

Data curation: Tsai-Min Chen, Chih-Han Huang.

Formal analysis: Tsai-Min Chen, Edward S. C. Shih.

Funding Acquisition: Ming-Jing Hwang.

Investigation: Tsai-Min Chen, Chih-Han Huang, Edward S. C. Shih.

Methodology: Tsai-Min Chen, Chih-Han Huang, Edward S. C. Shih, Ming-Jing Hwang.

Project Administration: Yu-Feng Hu, Ming-Jing Hwang.

Resources: Yu-Feng Hu, Tsai-Min Chen.

Software: Tsai-Min Chen, Chih-Han Huang, Edward S. C. Shih.

Supervision: Yu-Feng Hu, Ming-Jing Hwang.

Validation: Yu-Feng Hu, Chih-Han Huang, Edward S. C. Shih.

Visualization: Tsai-Min Chen, Edward S. C. Shih.

Writing – Original Draft Preparation: Tsai-Min Chen, Chih-Han Huang, Edward S. C. Shih, Yu-Feng Hu, Ming-Jing Hwang.

Writing – Review & Editing: Tsai-Min Chen, Chih-Han Huang, Edward S. C. Shih, Yu-Feng Hu, Ming-Jing Hwang.

## Supporting information captions

**S1 Table.** CPSC2018’s top 10 models and results (reported by the conference on http://2018.icbeb.org/Challenge.html)*

| Rank | Overall F1 | F<sub>af</sub> | F<sub>block</sub> | F<sub>pc</sub> | F<sub>st</sub> |
| --- | --- | --- | --- | --- | --- |
| **1** | **0.837** | **0.933** | **0.899** | **0.847** | **0.779** |
| 2 | 0.830 | 0.931 | 0.912 | 0.817 | 0.761 |
| 3 | 0.806 | 0.914 | 0.879 | 0.801 | 0.742 |
| 4 | 0.802 | 0.918 | 0.89 | 0.789 | 0.718 |
| 5 | 0.791 | 0.924 | 0.882 | 0.779 | 0.709 |
| 6 | 0.783 | 0.905 | 0.902 | 0.722 | 0.708 |
| 7 | 0.782 | 0.911 | 0.891 | 0.775 | 0.670 |
| 8 | 0.778 | 0.921 | 0.858 | 0.797 | 0.676 |
| 9 | 0.776 | 0.906 | 0.876 | 0.773 | 0.711 |
| 10 | 0.766 | 0.894 | 0.857 | 0.733 | 0.683 |

\*Our team's results are ranked first (boldfaced), and the highest scores of each sub-competition are indicated in red color. Overall F1 is the average of the F1 values from each classification type. F<sub>af</sub>: F1 of AF; F<sub>block</sub>: F1 of I-AVB, LBBB and RBBB; F<sub>pc</sub>: F1 of PAC and PVC; F<sub>st</sub>: F1 of STD and STE.

**S2 Table.** Comparisons on CA diagnosis between conference-assigned label, our model prediction, and consensus from three expert cardiologists.

| PatientID | Conf. | Model | Cardiologist1 | Cardiologist2 | Cardiologist3 | Consensus |
| --- | --- | --- | --- | --- | --- | --- |
| 6268 | STE | Normal | Normal | SR (normal) | SR (normal) | Normal |
| 6215 | STD | AF | narrow-QRS<br>tachycardia, SVT,<br>RBBB, RAD | PSVT | PSVT | PSVT |
| 4237 | RBBB | I-AVB | I-AVB | I-AVB, SR, TWI (V1-V4), rSR' (V1) | I-AVB,SR | I-AVB |
| 6380 | STE | LBBB | LBBB | LBBB,SR,LAE | LBBB,SR | LBBB |
| 1452 | AF | RBBB | AF, MVR, RBBB | AF, RBBB, Q wave, STE with<br>reciprocal change, w/o old MI | AF, RBBB | RBBB |
| 5398 | PVC | PAC | PAC | PAC | PAC,SR | PAC |
| 5963 | STD | PVC | PVC | PVC,SR,STD(V3-V6) | PVC,SR | PVC |
| 4278 | RBBB | STD | normal | SR, minimal STTC(II, III, aVF) | STD-like, SR | Normal |
| 504 | Normal | STE | STE-like | SR, early repolarization | SR | Normal |

Conference assignments (regarded as ‘ground truth’ for model training) or our model predictions in agreement with the consensus of the three expert cardiologists are highlighted in red. SVT: Supraventricular tachycardia; RAD: Right Axis Deviation; MVR: mitral valve
replacement; SR: sinus rhythm; PSVT: Paroxysmal supraventricular tachycardia; LAE: left atrial enlargement; MI: myocardial infarction; STTC: ST-T change

**S3 Table.** The performances of models trained without multi-labeled data*

| CA Type | Best Validation Models |  |  | Ensemble Model |  |  |
| --- | --- | --- | --- | --- | --- | --- |
|  | Median Accuracy | Median AUC (95% CI) | Median F1-score | Median Accuracy | Median AUC (95% CI) | Median F1-score |
| Normal | 0.940 | 0.908<br>(0.791-0.916) | 0.807 | 0.937 | 0.901<br>(0.808-0.932) | 0.794 |
| AF | 0.974 | 0.949<br>(0.885-0.992) | 0.915 | 0.980 | 0.955<br>(0.929-0.995) | 0.935 |
| I-AVB | 0.973 | 0.918<br>(0.852-0.991) | 0.876 | 0.976 | 0.912<br>(0.900-0.996) | 0.879 |
| LBBB | 0.993 | 0.927<br>(0.748-1.000) | 0.870 | 0.993 | 0.913<br>(0.770-1.000) | 0.862 |
| RBBB | 0.961 | 0.954<br>(0.895-0.981) | 0.922 | 0.944 | 0.925<br>(0.880-0.972) | 0.885 |
| PAC | 0.957 | 0.852<br>(0.747-0.961) | 0.747 | 0.965 | 0.889<br>(0.798-0.984) | 0.796 |
| PVC | 0.973 | 0.926<br>(0.832-0.989) | 0.858 | 0.974 | 0.950<br>(0.820-0.992) | 0.870 |
| STD | 0.950 | 0.869<br>(0.800-0.966) | 0.786 | 0.956 | 0.914<br>(0.790-0.950) | 0.821 |
| STE | 0.974 | 0.667<br>(0.491-0.995) | 0.394 | 0.975 | 0.663<br>(0.491-0.995) | 0.444 |

\* In these models, the 476 multi-labeled recordings were not included in the training set. These are results of the best validation models and the ensemble model on the ten 10-fold tests. These performances are comparable with those presented in Table 1.

**S4 Table.** Successfully predicted CA types for the 476 multi-labeled subjects in the released dataset of CPSC2018*

|  | AF | I-AVB | LBBB | RBBB | PAC | PVC | STD | STE |
| --- | --- | --- | --- | --- | --- | --- | --- | --- |
| AF | 0/0 | 0/0 | 17/29 | **154/172** | 0/4 | 6/8 | 4/33 | 0/2 |
| I.AVB |  | 0/0 | 2/8 | 8/10 | 0/3 | 2/5 | 0/6 | 0/4 |
| LBBB |  |  | 0/0 | 0/0 | 2/10 | 4/6 | 0/3 | 0/4 |
| RBBB |  |  |  | 0/0 | **34/55** | **49/51** | 16/20 | 5/19 |
| PAC |  |  |  |  | 0/0 | 3/3 | 4/6 | 2/5 |
| PVC |  |  |  |  |  | 0/0 | 6/18 | 0/2 |
| STD |  |  |  |  |  |  | 0/0 | 2/2 |
| STE |  |  |  |  |  |  |  | 0/0 |

\* Only the upper triangle portion of the symmetrical concurrent CA label counts is shown. The numbers shown are the number of correctly predicted subjects / the total multi-labeled subjects for a given CA type. The two CA types with the highest and the second highest probabilities are the predicted concurrent CA types. Boldfaced are the three most concurrent CA labels in these subjects (see Table 2).

**S5 Table.** The Bayes factors* (in log scale) of each lead’s performance (F1 score) relative to that of the best performing lead in each CA type (see Fig. 3)

|  | I | II | III | aVR | aVL | aVF | V1 | V2 | V3 | V4 | V5 | V6 |
| --- | --- | --- | --- | --- | --- | --- | --- | --- | --- | --- | --- | --- |
| Normal | 3.4 | 0.3 | 14.1 | -0.9 | 3.1 | 4.1 | 7.3 | 5.3 | 5.0 | 0.9 | 0.4 | -0.1 |
| AF | -0.8 | -0.8 | -0.6 | -0.9 | -0.3 | -0.7 | -0.9 | 0.2 | 1.1 | 0.8 | -0.3 | -0.2 |
| I-AVB | 1.2 | 0.1 | 3.7 | -0.4 | 0.1 | 0.8 | -0.9 | 1.3 | -0.5 | 0.5 | 1.1 | -0.1 |
| LBBB | -0.9 | 6.1 | 6.4 | -0.8 | 0.4 | 6.1 | -0.9 | 0.2 | 1.0 | 3.8 | 1.9 | -0.1 |
| RBBB | 18.3 | 15.5 | 22.2 | 11.4 | 4.1 | 18.8 | -0.9 | 9.9 | 12.9 | 16.0 | 18.6 | 1.8 |
| PAC | -0.9 | -0.9 | -0.6 | -0.3 | -0.1 | -0.9 | -0.9 | -0.8 | -0.7 | -0.2 | -0.4 | -0.2 |
| PVC | -0.8 | -0.9 | -0.8 | -0.8 | -0.6 | -0.9 | -0.9 | -0.9 | -0.9 | -0.9 | -0.9 | -0.6 |
| STD | 6.0 | -0.9 | 15.4 | -0.9 | 10.2 | 2.4 | 14.9 | 11.9 | 5.6 | 0.6 | -0.8 | -0.5 |
| STE | 11.9 | 1.7 | 11.3 | 3.7 | 23.1 | 6.7 | 12.2 | 7.1 | 0.3 | -0.9 | 0.4 | 0.8 |

\*computed using the 'BayesFactor' routine in the R package.

**S6 Table.** Leads in the top-performing group (threshold: Bayes factor<3.0)*

|  | I | II | III | aVR | aVL | aVF | V1 | V2 | V3 | V4 | V5 | V6 |
| --- | --- | --- | --- | --- | --- | --- | --- | --- | --- | --- | --- | --- |
| Normal | 0 | 1 | 0 | 1 | 0 | 0 | 0 | 0 | 0 | 0 | 1 | 1 |
| AF | 1 | 1 | 1 | 1 | 1 | 1 | 1 | 1 | 0 | 0 | 1 | 1 |
| I-AVB | 0 | 1 | 0 | 1 | 1 | 0 | 1 | 0 | 1 | 0 | 0 | 1 |
| LBBB | 1 | 0 | 0 | 1 | 1 | 0 | 1 | 1 | 0 | 0 | 0 | 1 |
| RBBB | 0 | 0 | 0 | 0 | 0 | 0 | 1 | 0 | 0 | 0 | 0 | 0 |
| PAC | 1 | 1 | 1 | 1 | 1 | 1 | 1 | 1 | 1 | 1 | 1 | 1 |
| PVC | 1 | 1 | 1 | 1 | 1 | 1 | 1 | 1 | 1 | 1 | 1 | 1 |
| STD | 0 | 1 | 0 | 1 | 0 | 0 | 0 | 0 | 0 | 0 | 1 | 1 |
| STE | 0 | 0 | 0 | 0 | 0 | 0 | 0 | 0 | 1 | 1 | 1 | 0 |
| Total | 4 | **6** | 3 | **7** | 5 | 3 | **6** | 4 | 4 | 3 | **6** | **7** |

\*The leads in the top-performing group, indicated by 1 (0 for those excluded from this group), for a given CA type are considered to perform equally well statistically based on the threshold of Bayes factor<3.0, which indicates the null hypothesis of no difference from the leading lead holds. Using this threshold, the sum total shows that leads aVR and V6 received most top-performing group counts, followed by leads II, V1, and V5.

**S7 Table.** Leads in the top-performing group (threshold: Bayes factor<0.33)*

|  | I | II | III | aVR | aVL | aVF | V1 | V2 | V3 | V4 | V5 | V6 |
| --- | --- | --- | --- | --- | --- | --- | --- | --- | --- | --- | --- | --- |
| Normal | 0 | 0 | 0 | 1 | 0 | 0 | 0 | 0 | 0 | 0 | 0 | 0 |
| AF | 1 | 1 | 1 | 1 | 0 | 1 | 1 | 0 | 0 | 0 | 0 | 0 |
| I-AVB | 0 | 0 | 0 | 0 | 0 | 0 | 1 | 0 | 1 | 0 | 0 | 0 |
| LBBB | 1 | 0 | 0 | 1 | 0 | 0 | 1 | 0 | 0 | 0 | 0 | 0 |
| RBBB | 0 | 0 | 0 | 0 | 0 | 0 | 1 | 0 | 0 | 0 | 0 | 0 |
| PAC | 1 | 1 | 1 | 0 | 0 | 1 | 1 | 1 | 1 | 0 | 0 | 0 |
| PVC | 1 | 1 | 1 | 1 | 1 | 1 | 1 | 1 | 1 | 1 | 1 | 1 |
| STD | 0 | 1 | 0 | 1 | 0 | 0 | 0 | 0 | 0 | 0 | 1 | 1 |
| STE | 0 | 0 | 0 | 0 | 0 | 0 | 0 | 0 | 0 | 1 | 0 | 0 |
| Total | 4 | 4 | 3 | **5** | 1 | 3 | **6** | 2 | 3 | 2 | 2 | 2 |

\*The leads in the top-performing group, indicated by 1 (0 for those excluded from this group), for a given CA type are considered to perform equally well statistically based on the threshold of Bayes factor<0.03, which indicates the null hypothesis of no difference from the leading lead holds. Using this threshold, the sum total shows that lead V1 received most top-performing group counts, followed by lead aVR.

## Notes

http://2018.icbeb.org/Challenge.html

